# No evidence for social genetic effects or genetic similarity among friends beyond that due to population stratification: a reappraisal of Domingue et al (2018)

**DOI:** 10.1101/643304

**Authors:** Loic Yengo, Morgan Sidari, Karin J. H. Verweij, Peter M. Visscher, Matthew C. Keller, Brendan P. Zietsch

## Abstract

Using data from 5,500 adolescents from the National Longitudinal Study of Adolescent to Adult Health, Domingue et al. (2018) claimed to show that friends are genetically more similar to one another than randomly selected peers, beyond the confounding effects of population stratification by ancestry. The authors also claimed to show ‘social-genetic’ effects, whereby individuals’ educational attainment (EA) is influenced by their friends’ genes. Neither claim is justified by the data. Mathematically we show that 1) although similarity at causal variants is expected under assortment, the genome-wide relationship between friends (and similarly between mates) is extremely small (an effect that could be explained by subtle population stratification) and 2) significant association between individuals’ EA and their friends’ polygenic score for EA is expected under homophily with no socio-genetic effects.

---

The availability of large samples of individuals with genome-wide genetic data in combination with behavioural phenotypes and social outcomes has led to a resurgence in research that address questions at the interface of genetics and the social sciences. Some of that research is hypothesis driven while much of it is data-driven and hypothesis generating. The genetics and statistical analysis of human traits has a solid underpinning theory in quantitative and population genetics (Lynch and Walsh 1998^1^; Walsh and Lynch 2018^2^), and rigorous benchmarking against these underpinnings is essential, especially when novel or unexpected results in human behaviour are reported. In this short note, we highlight one example (and list others) where novel results and claims are not justified by the data presented and instead have alternative and more parsimonious explanations. Using data from 5,500 adolescents from the National Longitudinal Study of Adolescent to Adult Health, Domingue et al.^3^ claimed to show that friends are genetically more similar to one another than randomly selected peers, beyond the confounding effects of population stratification by ancestry. The authors also report evidence of ‘social-genetic’ effects, whereby individuals’ educational attainment (EA) is influenced by their friends’ genes. Here we argue that neither claim is justified by the data.

First, we show that individuals who match on phenotype – whether as romantic partners or, in this case, friends – do not exhibit a substantially increased genome-wide similarity. Indeed, Robinson et al.^4^ previously showed that the expected genomic relatedness between individuals phenotypically matching on a given trait equals *rh*^*2*^*/M*_*e*_, where *r* is the phenotypic correlation between matched individuals, *h*^2^ the heritability of the trait driving the assortment and *M*_*e*_ the effective number of independent markers in the genome, estimated in European descents ∼50,000.^5^ Therefore, assuming the SNP heritability of EA to be *h*^2^∼0.12 (Ref.^6^) and given the correlation of EA between friends (*r*=0.415; from their Table 2) reported by Domingue et al., we would expect a genomic relatedness between friends of 0.12×0.415/50,000, i.e. ∼10^-6^. Unfortunately, the friend-pair genomic similarity of 0.031 (95% confidence interval (CI): 0.022 – 0.036) reported by Domingue et al.^3^ does not represent an actual estimate of genomic relationship as classically used in the human genetics literature^7^, but is instead an alternative measure introduced by the authors in a previous publication^8^.

In **Supplementary Note 1**, we derive the mathematical relation between Domingue et al.’s similarity measure and the classical genomic relationship. We show that a similarity of 0.031 on the authors’ scale is akin to a genomic relationship between friends of ∼5×10_-4_. Although quite small, this value is still over two orders of magnitude larger than the theoretical value of ∼10^-6^ expected under phenotypic matching. Given that phenotypic matching cannot explain this large discrepancy, the most likely cause of such genome-wide similarity among friends would be population stratification, whereby individuals are more likely to befriend others living in close geographical vicinity and thus of likely similar ancestry. Domingue et al. acknowledged the possibility of confounding due to population stratification, but in our view did not control for it in a standard way. Rather, they reported (in their Supplementary Materials) a secondary analysis using the program REAP, which they argue is robust to stratification and still reveals a significant (though reduced: from 0.031 to 0.02 (95%CI: 0.011 – 0.028); data from Table S3 in ref.^3^) genomic similarity among friends. However, REAP was designed to estimate kinship among related individuals in admixed samples with heterogeneous continental ancestry^9^ – not for estimating genomic similarity among unrelated individuals with homogenous continental ancestry (e.g. Domingue et al.’s sample of European ancestry; T. Thornton, personal communication, April 8, 2018). Given these limitations, it appears to us the most likely and more parsimonious explanation for the observed genomic similarity among friends is within-continental population stratification (e.g. Northern vs. Southern European ancestry).

Second, Domingue et al.’s results do not provide evidence for ‘social-genetic’ effects as claimed. The purported evidence is a significant association between focal individuals’ EA and their friends’ polygenic score for EA (PGS_EA_), controlling for focal individuals’ PGS_EA_. We show mathematically in **Supplementary Note 2** that simple homophily – the well-established tendency for individuals to befriend others with similar educational performance^10,11^ – can explain the significant association that Domingue et al. used as evidence for a complex socio-genetic process. While our analyses do not rule out the possibility of ‘social-genetic effects’, other research using longitudinal sample of high school and university students (*N*=6,000), showed that friend similarity in academic performance was due to initial choice of similar friends, and not to a change in individuals’ academic performance towards that of their friends^10^. This finding is inconsistent with ‘social-genetic effects’ as envisaged by Domingue et al.

In summary, the advent of large samples of genotyped individuals with known social relationships has provided unprecedented opportunities for research at the intersection of human genetics and social sciences. However, analysis and interpretation of these data require great care, and several high-profile papers^8,12^ on the genetic similarity of social or romantic mates have forwarded exciting but perhaps misleading interpretations of results that probably have more parsimonious explanations, such as population stratification (e.g. see commentaries refs.^13,14^ on papers refs.^8,12^).

## Acknowledgements

This research was supported by the Australian Research Council (DP160102400; FT160100298), the Australian National Health and Medical Research Council (1113400 and 1078037) and the National Institute of Health (NIH grants R01AH042568 and R01MH100141). K.J.H.V. is supported by the Foundation Volksbond Rotterdam.

## Supplementary Notes

### Note 1: Interpretation of Domingue et al.’s genetic similarity in terms of kinship differences

In Domingue et al.^8^, the authors introduced a new metric to quantify the similarity within pairs of a certain class (e.g. friends or mates) relative to the similarity of random pairs. That new statistic, hereafter denoted *I*(*µ*), is an estimator of the area under the curve defined by the quantiles of the distribution of kinship coefficients under the null (random mating) versus the quantiles of the distribution of kinship coefficients under the alternative (e.g. assortative mating). We derive below an interpretation of that statistic in terms of differences in mean kinship between the two groups of pairs.

We consider two distributions of kinship coefficients: under the null (**H**_0_: *N*(0,*σ*^2^)) and under the alternative (**H**_1_: *N*(−*µ,σ*^2^)). The area of the shaded zone (Fig 1. in Domingue et al.^8^), hereafter denoted *I*(*µ*), is therefore defined as

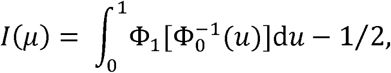

where Φ_*k*_ is the cumulative distribution function of relatedness coefficients under **H**_k_.

If we posit 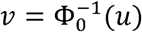, i.e. *u* = Φ_0_(*v*), then d*u* = *ϕ*_0_(*v*)d*v*, with *ϕ*_0_(.) being the probability density function of relatedness under the null. When 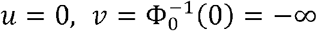 and when 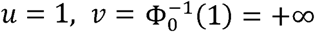. Therefore, *I*(*µ*) can rewritten as

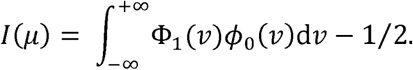

Under Gaussian assumptions, we can show that Φ_l_(*v*) = Φ_0_(*v* + *µ*). Therefore,

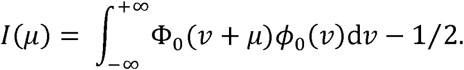

*I*(*µ*) cannot be calculated analytically. However, for small values of *µ* (near 0) we can derive its Taylor’s series as

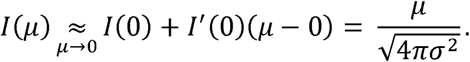

In Domingue et al. (2018), the reported genetic similarity between friends is .031 (Table 1). This implies a difference of kinship coefficients (0.5× the off-diagonal terms of Genetic Relationship Matrix, GRM), in friend pairs compared to random pairs, of ∼. 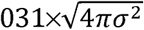. The variance of the off-diagonal terms of the GRM can be approximated as *σ*^2^ *≈* 1/*M*_*e*_ 1/50,000 = 2×10^-5^. (Visscher et al.^5^). Therefore a genetic similarity of .031 would correspond to a kinship difference of ∼2×10^-4^, i.e. a difference in GRM off-diagonal elements ∼0.0005. Such a small difference is more likely created by population stratification.

### Note 2: Mathematical expectations of ‘social genetic’ effects reported in Domingue et al

Domingue et al. define social-genetic effects as “the influence of one organism’s genotype on a different organism’s phenotype” (p. 704)—in this case, the effects of friends’ genomes on a focal individual’s phenotype. The purpose of this section is to argue that the results reported by Domingue et al. in support of social genetic effects among friends are more simply and parsimoniously explained by a model of individuals befriending others with similar potential for Educational Attainment (EA) values. Such a process is called homophily or “phenotypic assortment”. Such phenotypic assortment among friends on EA, combined with the fact that EA can be predicted from its polygenic score (PGS), necessarily implies correlations between friends’ PGS values as well as correlations between focal individuals’ EA values and their friends’ PGS values. While social-genetic effects cannot be ruled out, we show here that the relationships reported by Domingue et al. are expected consequences of simple assortment among friends on EA, and therefore there is no need to resort to more complicated explanations such as social-genetic effects.

We focus first on the central finding that Domingue et al. argue is evidence for social genetic-effects of EA: the EA of focal individuals is significantly related with the PGS of their friends. We derive this expected relationship based only on the EA-PGS relationship and the relationship between EAs of friends. We define the following terms:

- *EA*_*i*_: the EA of focal individual *i*
- 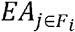: the EA of individual *j*, a member of individual *i*’s friend group (*F*_*i*_)
- *PGS*_*i*_: the PGS for EA of focal individual *i*
- 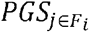: the PGS of EA of individual *j*, a member of individual *i*’s friend group (*F*_*i*_)
- 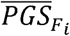: the mean PGS of EA across all friends of individual *i* (*F*_*i*_)
- 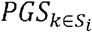: the PGS of EA of individual *k*, a member of individual *i*’s schoolmates (*S*_*i*_)
- 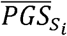: the mean PGS of EA across all schoolmates of individual *i* (*S*_*i*_)

Using path tracing rules in Figure RS1 below and the correlations presented in Table 2 (of the Domingue et al. manuscript), the expected correlation between *EA*_*i*_ and 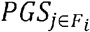 (or equivalently between *PGS*_*i*_ and 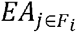) is .26 × .42 = 109, and the expected correlation between *PGS*_*i*_ and 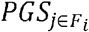 is .26 × .42 × .26 = .029 (all variables are standardized as noted on p. S3 of Domingue et al. Supplementary Materials).

**Figure RS1.**
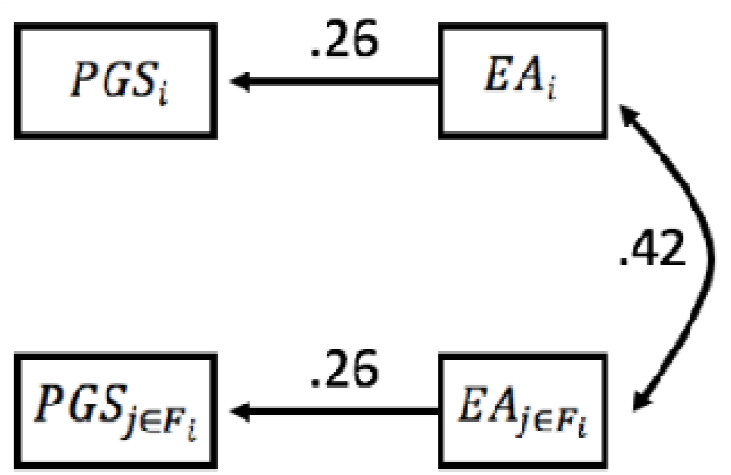

**Figure RS2.**
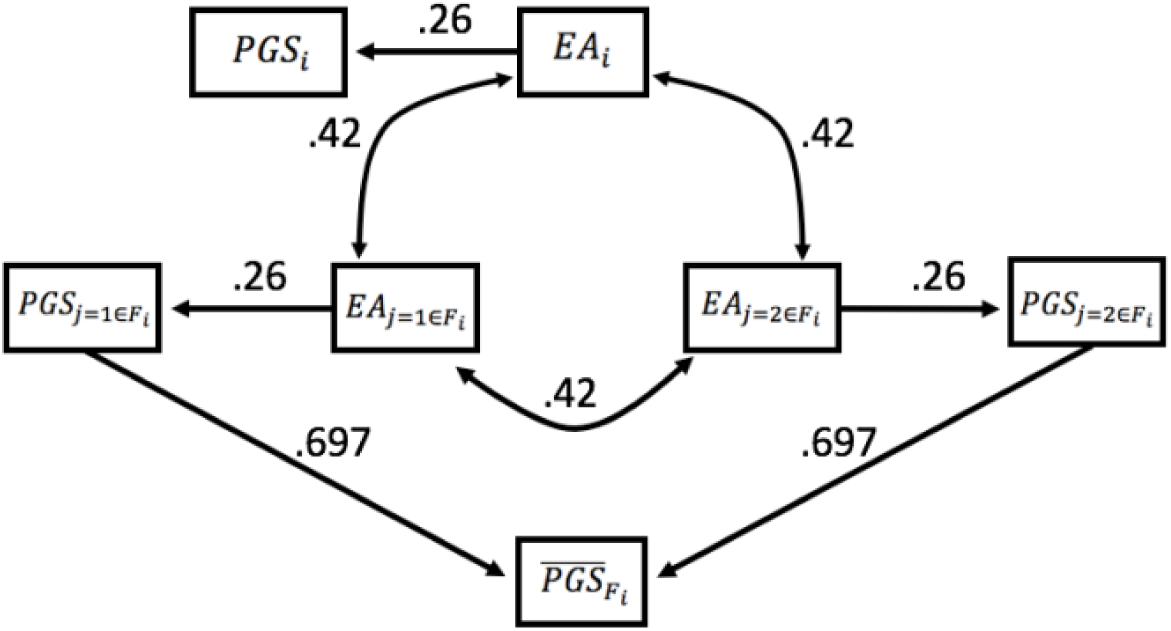

In their Figure 2 and Table S6, Domingue et al. report the relationship (the slope in this case) between and, which is different from the relationship between and derived above and depends on the number of friends included in the average *PGS* score,. The number of friends an individual had in Add Health varied across individual but had a mean of 2 (Figure S3). For mathematical tractability, we assume the number of friends was constant at 2 across all focal individuals. Furthermore, the slope of ∼ depends crucially on the variances of both variables. The authors state that outcomes and predictors were standardized for this analysis (caption, Figure 2), and our expectations below agree with this. We therefore assume that was standardized after taking the mean. The coefficients to in Figure RS2 below (.697) are those that lead to, after accounting for the correlation between and. Our model assumes that co-friends of a focal individual are correlated as highly as each friend is to the focal individual, but this assumption has only a minor influence on results: the coefficient from to would be only slightly different (^—^) if co-friends’ EA values were uncorrelated.

The expected slope of regressed on, can be expressed as ——— given that all variables are standardized. Using path tracing rules,, which agrees closely with the reported (Table S6, column 4).

Domingue et al. then control for and find that this partial slope is only slightly reduced () and still significant. They interpret this partial slope as evidence “…that the genetics of individuals in a person’s social environment influence that person’s phenotype,” (p. 705). However, controlling for a variable that is only weakly associated with the outcome and predictor variables, such as, is expected to change the slope by only a small amount. In particular, given that all variables are standardized,

Using path tracing rules and Figure RS2, the correlation between *PGS*_*i*_ and 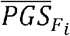 is 2 × [.697 × .26×.42 × .26] = .0396 ≅ .04. Thus, under the assumption only of phenotypic assortment among friends’ EA values, this expected partial slope is

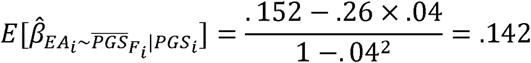

which is, again, close to the partial slope (.154 ± .03) reported in the manuscript. Similarly, the other relationships reported in Domingue et al. do not differ from what is expected under phenotypic assortment of EA. For example, the partial slope 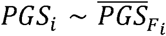 is expected to be .052 (vs. .06 ± .03 reported), and the partial slope, 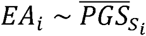 is expected to be .197 (vs. .22 .± 03 reported). (It should be noted that the results reported in Figure 1 and Table S4 appear to be regression coefficients where the predictors, 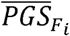 or 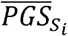, were not standardized after taking the mean. This is seen most clearly in the slope associated with 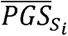 in Table S4 (.62), which is far too high to be a slope between the standardized values of these two variables. Our expectations assumed that 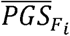 and 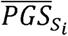 were unstandardized in these analyses).

In summary, the results interpreted by Domingue et al. as evidence for social-genetic effects are expected under a simple model of people befriending individuals of similar educational attainment values. The fact that individuals’ EA scores are correlated with their friends’ PGS scores is a necessary consequence of such assortment on EA. While more complicated models explaining these relationships are possible, they are not necessary to explain the presented results.

